# Precision mapping of functional brain network trajectories during early development

**DOI:** 10.1101/2025.06.16.659959

**Authors:** Diego Derman, Silvina L. Ferradal

## Abstract

Preterm birth is a known risk factor for neurodevelopmental disabilities, but early cognitive assessments often fail to predict long-term outcomes. This limitation underscores the need for alternative biomarkers that reflect early brain organization. Resting-state functional connectivity is a powerful tool to study functional brain organization during the perinatal period. However, most fMRI studies in infant populations use group-level analyses that average subject-specific data across several weeks of development, reducing sensitivity to subtle, time-sensitive deviations from typical brain trajectories. Using a novel precision functional mapping approach, we estimated individual resting-state networks (RSNs) in a large cohort of neonates (N = 352, gestational age at birth: 25.6–42.3 weeks) from the developing Human Connectome Project. RSN connectivity strength increased linearly with age at scan, especially in higher-order networks. In particular, the default mode network (DMN) exhibited marked changes in topography and connectivity strength, evolving from an immature organization in preterm infants to a more adult-like pattern in term-born infants. Longitudinal data from a subset of preterm infants (N = 15) confirmed ongoing network development shortly after birth. Despite this maturation, preterm infants did not reach the connectivity levels of term-born infants by term-equivalent age. These findings highlight the potential of individualized RSN mapping as an early marker of neurodevelopmental trajectories.

## Introduction

Preterm birth is a well-established risk factor for neurodevelopmental disabilities later in life (Bhutta et al., 2002; Hadaya et al., 2025; Marlow et al., 2005; Saigal & Doyle, 2008). Yet, cognitive assessments administered in the first years of life remain poor predictors of long-term outcomes (Spencer-Smith et al., 2015; Spittle et al., 2015). Functional brain mapping has the potential to reveal meaningful relationships between early brain organization and later behavioral outcomes (Bijsterbosch et al., 2018; Cui et al., 2020; Finn et al., 2015; Kong et al., 2021; Rosenberg et al., 2016). Resting-state functional connectivity measured with functional magnetic resonance imaging (fMRI) has enabled the identification of resting-state networks (RSNs), providing insights into the early functional organization of the infant brain (Fransson et al., 2011, 2009, 2007; Gao et al., 2009). Previous studies have reported both qualitative and quantitative differences in RSNs between term and preterm-born infants (Cao et al., 2016; Doria et al., 2010; Smyser et al., 2010), as well as early associations between RSNs and developmental phenotypes such as age (Eyre et al., 2021). However, most existing work has emphasized group-level analyses, potentially hiding meaningful individual variability (Labonte et al., 2024).

While traditional group-level analyses have revealed shared features of functional organization (Gordon et al., 2016), recent work in adults has shown that these approaches mask individual-specific patterns (Gratton et al., 2020), limiting their ability to predict behavioral differences (Sylvester et al., 2020). Given that both prenatal and postnatal experiences play a critical role in shaping functional brain development (Labonte et al., 2024), individualized analyses of functional connectivity during the perinatal period may provide valuable insights into early, personalized developmental trajectories.

Despite its promise, estimating individual-level RSNs in neonates remains challenging due to the unique constraints of fMRI data in young infants. In this study, we apply a novel precision functional mapping approach (Derman et al., 2025) to estimate RSNs in individual neonates using limited fMRI data. Leveraging the large dataset from the developing Human Connectome Project (Edwards et al., 2022), we built cross-sectional maturation curves in a cohort of 352 preterm and term-born infants (gestational age at birth: 25.6 – 42.3 weeks). We found a significant linear association between age at scan and connectivity strength across all RSNs. Moreover, preterm birth was associated with a significant reduction in connectivity strength across most networks. These findings were reinforced by a subset of preterm infants (N = 15) who underwent two scans, one at preterm age and another at term-equivalent age, revealing clear evidence of RSN maturation over time, particularly in higher-order networks. Notably, our results suggest that the default mode network (DMN) undergoes a developmental transition during the perinatal period, evolving from an immature state to one that more closely resembles its adult organization.

Overall, these findings demonstrate the potential of precision functional mapping to capture individual differences in early brain organization. This approach holds promise for advancing our understanding of early functional development and may ultimately aid in the early identification of at-risk infants and the design of personalized interventions.

## Methods

### Subjects and MRI data acquisition

MRI data was obtained from the second release of the developing Human Connectome Project (dHCP) database. For this study, we included subjects with complete functional imaging sessions that passed quality control and had a radiological score of two or lower, indicating no lesions of clinical or analytical significance. Based on these criteria, 391 sessions corresponding to 371 subjects (age at birth: 25.6 – 42.3 weeks gestational age, GA) scanned shortly after birth or at term-equivalent age (age at scan: 29.3 – 44.9 weeks postmenstrual age, PMA) were considered for further analysis.

All scans were obtained with a 3T Philips Achieva using a neonatal head coil at Evelina Newborn Imaging Centre, St. Thomas Hospital, London, UK. Both T1-weighted (TR = 4795 ms; TE = 8.7 ms) and T2-weighted (TR = 12 s; TE = 156 ms) structural scans were obtained with a multi-slice Turbo Spin Echo (TSE) sequence, with in-plane resolution 0.8 x 0.8 mm² and 1.6 mm slices overlapped by 0.8 mm. Two stacks of images were taken per weighting, sagittal and axial, which were integrated to obtain T2w volumes with an image resolution of 0.8 mm isotropic. Blood oxygen level-dependent (BOLD) scans were obtained with a multi-slice gradient-echo planar imaging (EPI) sequence (TE = 38 ms; TR = 392 ms, multiband factor = 9; flip angle = 34°) with an image resolution of 2.15 mm isotropic. A resting-state BOLD fMRI acquisition of 2300 time points (15 minutes) was obtained for each infant.

### MRI Preprocessing

Structural and functional MRI data were preprocessed as described in Derman et al., 2025. Briefly, individual cortical surfaces from the dHCP minimal processing pipeline (Makropoulos et al., 2018) were aligned to a 40-week symmetrical atlas (Williams et al., 2023) using the Multimodal Surface Matching tool based on cortical folding (Robinson et al., 2018). Resting-state BOLD fMRI data from the preprocessed dHCP pipeline (Fitzgibbon et al., 2020) were projected onto each individual surface using a modified version of the HCP surface pipeline (Glasser et al., 2013) and subsequently aligned to the cohort-specific 40-week atlas. Geodesic 2D Gaussian smoothing was applied using a 3 mm FWHM kernel.

Following a conservative frame censoring approach, a contiguous block of 1600 frames (10 minutes) with the minimum number of motion outliers (as measured by DVARS) was retained for each subject. Sessions with more than 160 motion-corrupted volumes (10%) were excluded. Based on this criterion, 369 sessions from 352 subjects (274 term-born and 78 preterm infants) were included in the analysis. Table 1 presents a summary of the demographics considered in the analysis.

**Table 1:**
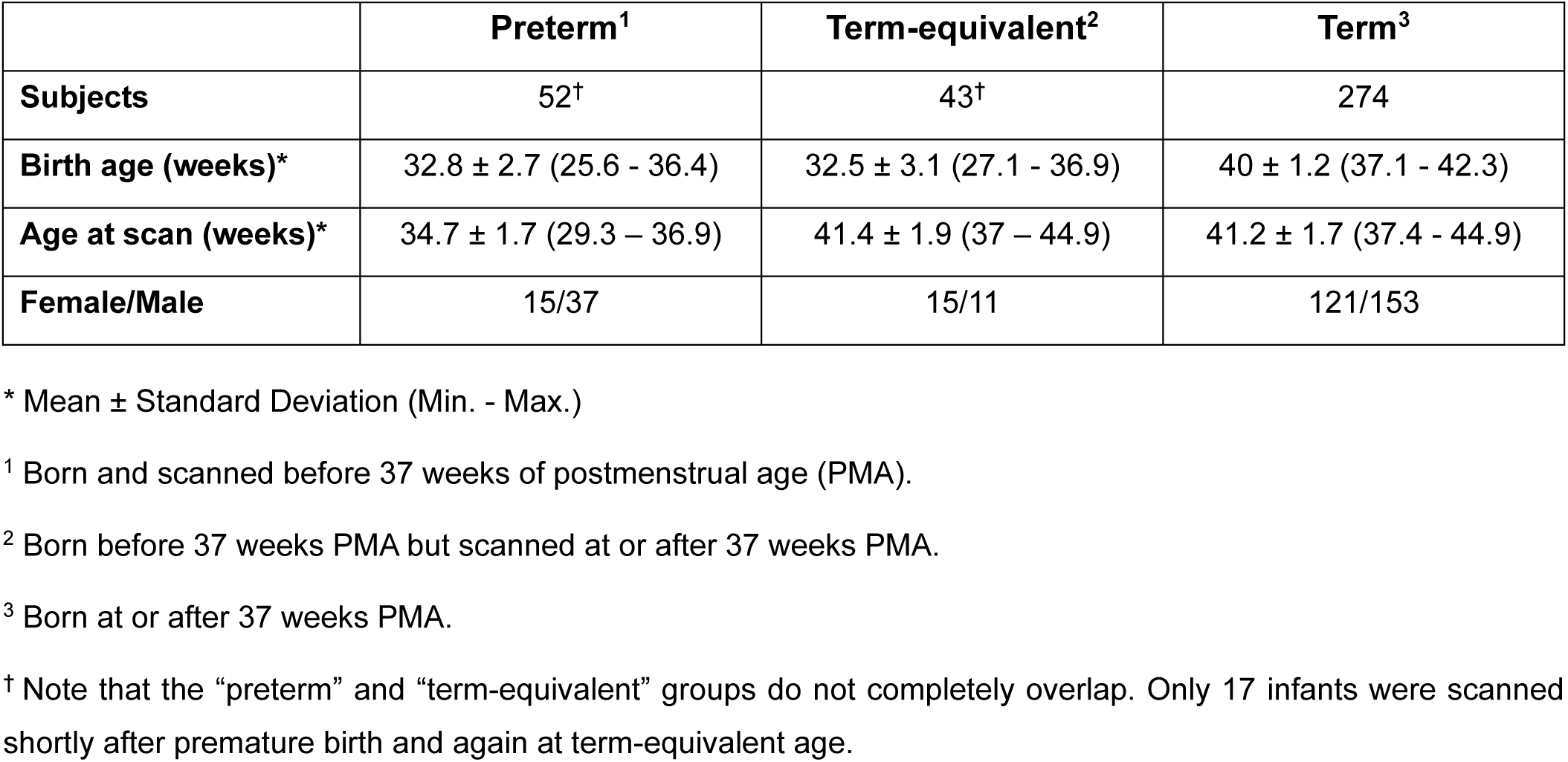
Study population. After excluding subjects with excessive motion artifacts, 352 infants with 369 scans from the second release of the dHCP dataset were included in the present study. Participants were categorized into three groups - preterm, term-equivalent, and term - based on their birth age and age at scan.

|  | Preterm<sup>1</sup> | Term-equivalent<sup>2</sup> | Term<sup>3</sup> |
| --- | --- | --- | --- |
| **Subjects** | 52<sup>†</sup> | 43<sup>†</sup> | 274 |
| **Birth age (weeks)\*** | 32.8 ± 2.7 (25.6 - 36.4) | 32.5 ± 3.1 (27.1 - 36.9) | 40 ± 1.2 (37.1 - 42.3) |
| **Age at scan (weeks)\*** | 34.7 ± 1.7 (29.3 – 36.9) | 41.4 ± 1.9 (37 – 44.9) | 41.2 ± 1.7 (37.4 - 44.9) |
| **Female/Male** | 15/37 | 15/11 | 121/153 |

\* Mean ± Standard Deviation (Min. - Max.)
<sup>1</sup> Born and scanned before 37 weeks of postmenstrual age (PMA).
<sup>2</sup> Born before 37 weeks PMA but scanned at or after 37 weeks PMA.
<sup>3</sup> Born at or after 37 weeks PMA.
<sup>†</sup> Note that the “preterm” and “term-equivalent” groups do not completely overlap. Only 17 infants were scanned shortly after premature birth and again at term-equivalent age.

### Subject-level estimation of RSN maps

To obtain individual resting-state network (RSN) maps, we used a Bayesian hierarchical model that incorporates population information through empirical priors (Mejia et al., 2020). For this study, the empirical priors include estimates of the mean and between-subject variance of RSNs from a representative sample of 36 infants uniformly distributed across the entire cohort (age at scan: 36.6 – 44.9 weeks PMA, birth age: 29.4 – 41.9 weeks PMA). Note that the subjects included in the empirical priors were removed from further analysis.

To construct these priors, we first performed group independent component analysis (ICA) on a subset of 24 infants (age at scan: 44.5 – 45.5 weeks PMA). Based on visual inspection of the group ICA maps, associated time series, and power spectra, we identified eight RSN maps: lateral motor, medial motor, somatosensory, auditory, primary visual, motor association, visual association, and default mode network. We then applied dual regression to the 36-subject representative sample to obtain rough estimates of individual RSN maps, which were then used to compute the mean and the between-subject variance of each of the eight RSNs.

Using the empirical priors along with subject-specific fMRI data as inputs to an expectation-maximization (EM) algorithm, we obtained subject-level estimates of the posterior mean and standard deviation for each individual RSN map. Finally, we computed t-statistic maps from the subject-specific posterior mean and standard deviation maps. For a detailed description of the Bayesian framework, see our methodological paper (Derman et al., 2025).

## Results

Individual-level functional parcellations reveal system-level differences between preterm and term infants, as illustrated by representative subjects from the full cohort (Fig. 1). Note the stability of the primary sensory RSNs, such as medial and lateral motor networks. In contrast, higher-order RSNs seem to be still developing, as evidenced by the prefrontal and temporal clusters of the default mode network (DMN) and the motor association network.

**Figure 1:**
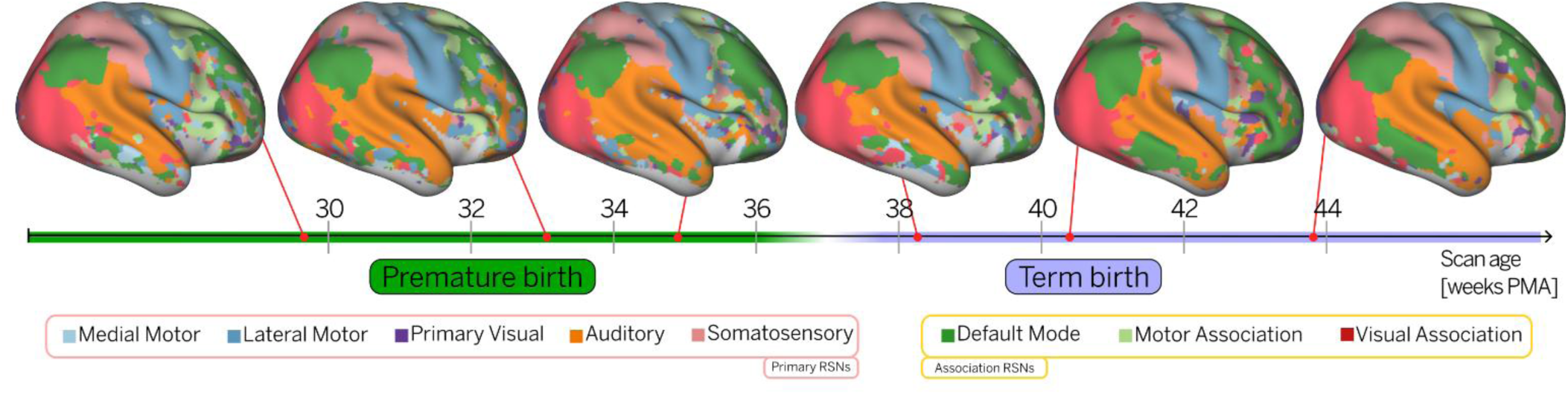
Individual-level parcellations reveal system-level differences in preterm and term-born infants. Functional parcellations for six randomly selected subjects scanned at different ages. The position of each subject on the timeline reflects their postmenstrual age (PMA) at the time of scanning (29.9, 32.6, 35.1, 38.1, 40.1, and 43.9 weeks PMA, respectively). Subjects born at or after 37 weeks PMA are considered term-born. Functional parcellations were obtained using a winner-takes-all (WTA) approach, where each vertex was assigned to the resting-state network (RSN) with the highest value at that location. The results were projected onto a 40-week cortical surface atlas for visualization purposes.

### Longitudinal trajectories of RSNs in premature infants

To characterize the maturation of RSNs in premature infants, we analyzed longitudinal changes in connectivity strength in a cohort of 15 infants scanned at two time points: once shortly after birth during the preterm period, and again at term-equivalent age. Note that two preterm infants from this longitudinal sample were excluded from further analysis because their data were used to construct the empirical priors. To quantify these changes, we first generated binary masks for each individual RSN by thresholding the t-statistic maps to identify regions of significant activation (see Methods). Then, we calculated mean connectivity strength for each individual RSN as the average t-statistic value within each mask, weighted by the surface area of each vertex. To minimize potential bias from motion differences between scans, we regressed out motion effects using a linear model. The model confirmed no significant differences in motion between the two sessions. Finally, we used a linear regression model to estimate the average maturation between the first and second scans. Across time, we observed increases in functional connectivity strength in all RSNs (Fig. 2), consistent with ongoing network maturation during this critical developmental window.

**Figure 2:**
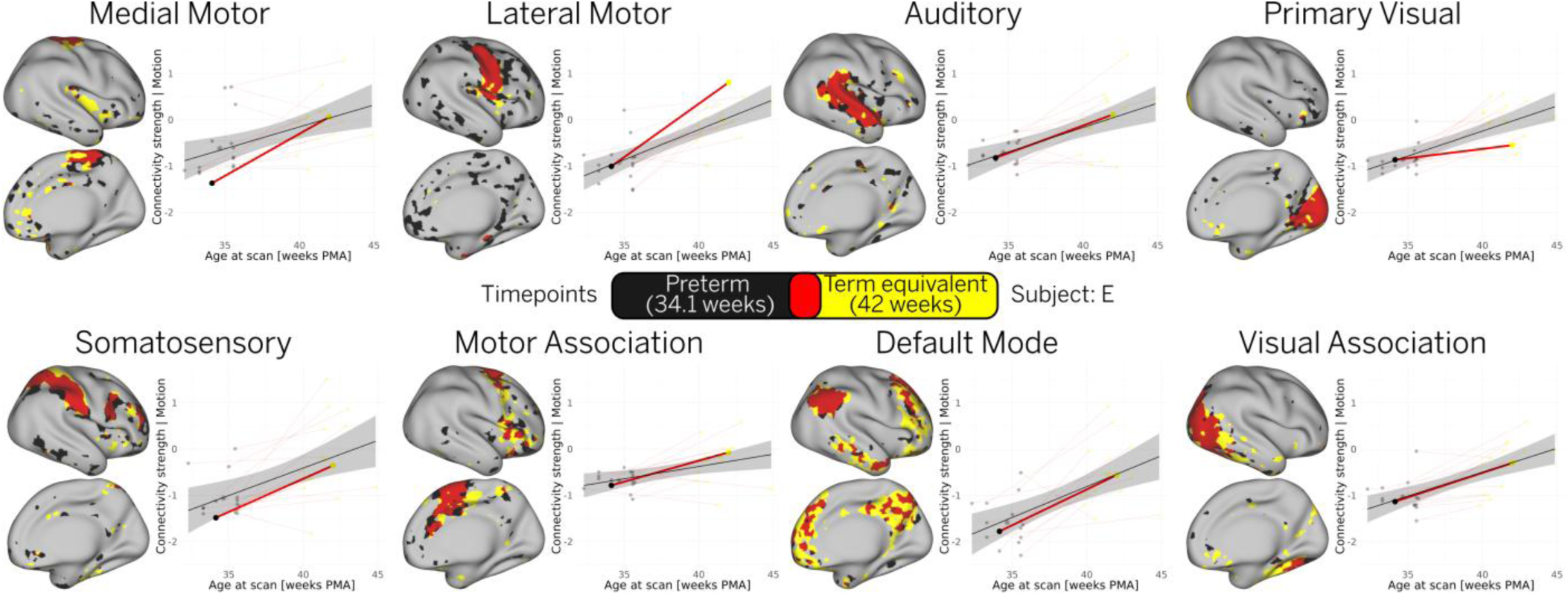
**Longitudinal trajectories in premature infants show increased connectivity strength across networks**. The relationship between age at scan and individual connectivity strength was examined in 15 preterm infants, each scanned twice: once at preterm age and again at term-equivalent age. The solid black line represents the group-level linear trend, with the shaded gray area indicating the 95% confidence interval. For illustration, we highlight one representative infant from this cohort, showing the individual linear trend (red) across the two scanning sessions, along with the associated functional parcellation maps for each RSN. In these maps, black, yellow, and red regions indicate parcels present at the first session (34.1 weeks PMA), the second session (42 weeks PMA), and their spatial overlap, respectively.

We also observed topographical changes in some RSNs between the two scans, as illustrated by individual-level functional parcellation maps from a representative infant in this cohort (Fig. 2). Notably, the primary sensory RSNs remained relatively stable, while topographical expansion was observed in the association RSNs, particularly the DMN.

### Trajectories of RSNs at term-equivalent age

Next, we examined whether the developmental trajectories of preterm infants converged with those of term-born infants by term-equivalent age. To investigate this, we compared the spatial topography and connectivity strength across all RSNs between preterm infants scanned at term-equivalent age (N = 37; 37–44.9 weeks PMA) and term-born infants scanned within the same age range (N = 247). Note that six preterm infants from the term-equivalent group and 27 from the term-born group were excluded from further analysis because their data were used to construct the empirical priors. Qualitative comparisons of group-average t-statistic maps revealed weaker connectivity patterns across all networks in preterm infants at term-equivalent age compared to term-born infants. Furthermore, a two-sided t-test of connectivity strength, adjusted for sex, motion, and scan age, revealed statistically significant differences across all networks except the medial motor network (Fig. 3).

**Figure 3:**
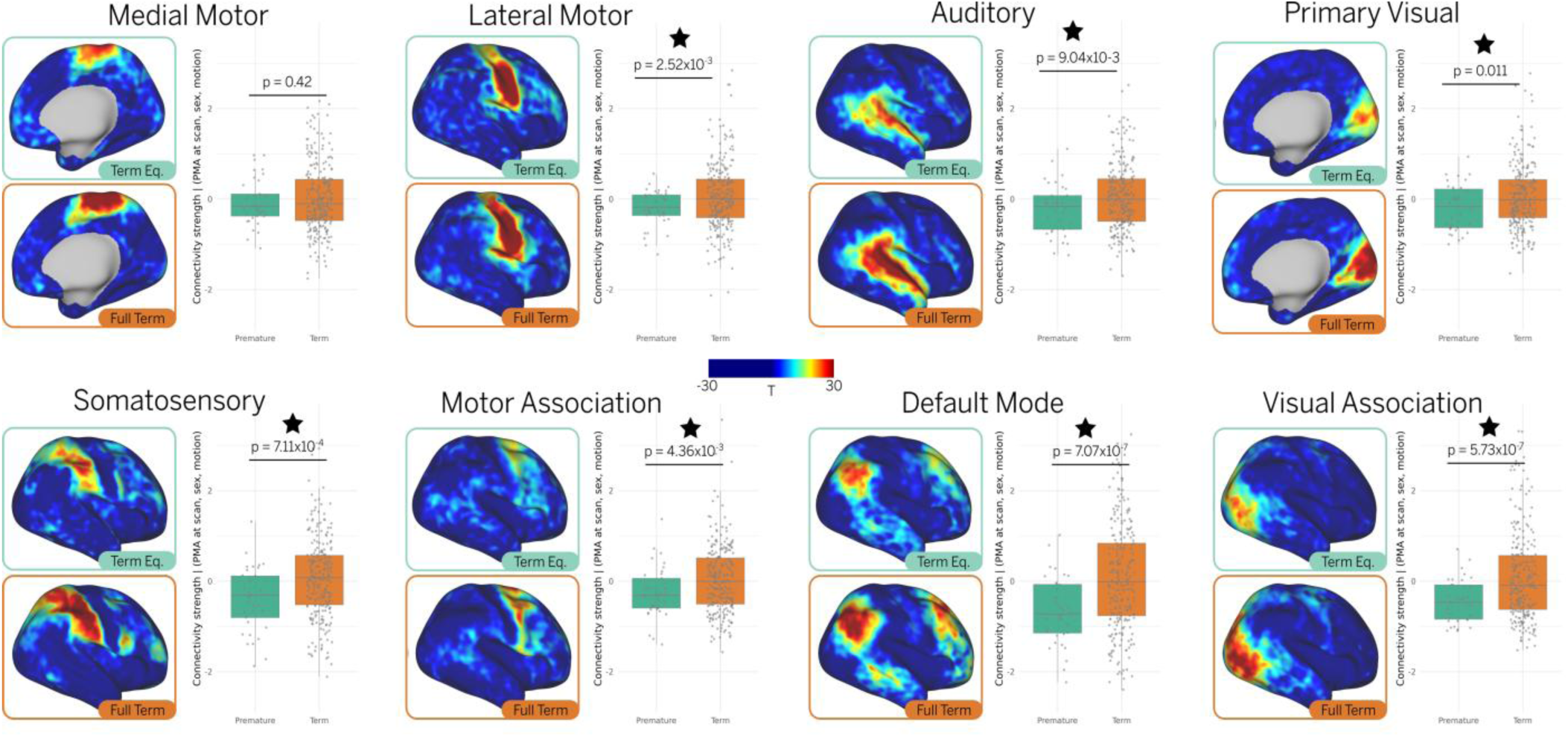
Functional connectivity profiles show differences between preterm and term- born infants at term-equivalent age. Group-average t-statistic maps for term-born (N = 247) and preterm infants scanned at term-equivalent age (N = 37) reveal generally weaker RSNs in preterm infants. This finding is supported by t-test comparisons, which show significantly lower connectivity strength in preterm infants across all networks except the medial motor network. For visualization, group maps are projected onto the 40-week inflated atlas. Stars represent statistically significant differences between groups (p<0.05).

### Effect of age and prematurity

Finally, we aimed to characterize the entire developmental trajectory of RSNs in term- and preterm-born infants during the early postnatal period. For each session in the cohort, we computed connectivity strength from individual-level RSN maps estimated with our Bayesian model and fit a linear regression with connectivity strength as a function of scan age. The regression model controlled for sex and motion and included an interaction term for prematurity to assess its influence on RSN maturation. A statistically significant relationship between connectivity strength and age at scan was found for all RSNs (Fig. 4). Furthermore, preterm infants show a significantly lower connectivity strength than term-born infants for all networks except medial motor. No significant interaction effect between prematurity and age at scan was observed, confirming that the developmental trajectories of preterm infants do not converge with those of the term-born infants within the studied age range.

**Figure 4.**
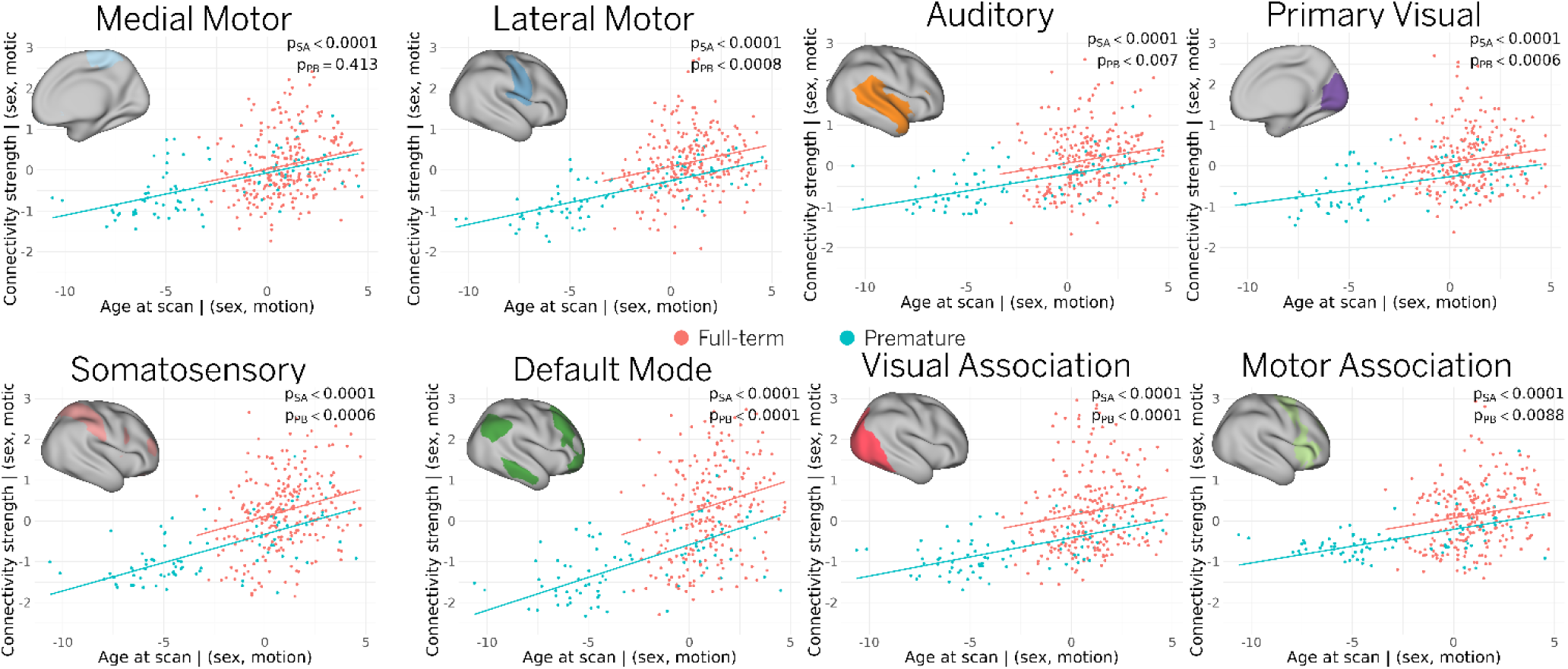
Effect of age and prematurity. A linear regression model was used to assess the relationship between individual connectivity strength and scan age across the entire cohort, controlling for sex and motion. All RSNs exhibited significant age-related increases in connectivity strength. However, preterm infants showed significantly lower connectivity than term-born infants in every network except the medial motor (as denoted by ppb). This offset persisted across the studied age range, with no difference in maturation rate between preterm and term-born groups. Note that pSA refers to the p-value for the linear term in the regression model, which captures the association between connectivity strength and age at scan (i.e., maturation). A value below 0.05 indicates a significant positive association in term-born infants, suggesting that connectivity increases with age. Similarly, ppb refers to the p-value for the categorical variable “preterm birth.” A value below 0.05 indicates that preterm infants exhibit significantly lower connectivity strength compared to term-born infants. For all RSNs, the p-values for the interaction term between “preterm birth” and “age at scan” were greater than 0.05, indicating that the relationship between connectivity and age does not significantly differ between term and preterm infants (i.e., both groups show a similar maturation pattern over time).

## Discussion

This study advances our understanding of how premature birth affects functional brain development by extending previous findings and introducing novel methodological improvements. Consistent with prior research, we observed weaker functional connectivity in preterm infants at term-equivalent age compared to term-born neonates (Eyre et al., 2021; Smyser et al., 2010; Sun et al., 2024). Our results also capture the ongoing maturation of resting-state networks (RSNs) during the early postnatal period, with significant increases in functional connectivity strength observed across all cortical networks.

### Individual-level precision mapping of functional maturation

A central contribution of this work is the ability to examine individual-level RSN topography in preterm and term-born neonates, enabling a more detailed analysis of early brain development. This was made possible by a precision functional mapping approach (Derman et al., 2025), which combines surface-based alignment, age-matched atlases, and Bayesian modeling to produce cleaner subject-level RSN estimates. These improvements allowed us to track topographical changes in individual RSNs at different ages (Fig. 1), which are not possible to capture using traditional group-level approaches. Our results suggest that higher-order RSNs undergo accelerated maturation following premature birth (Fig. 2). These networks also exhibit the largest differences in connectivity strength between preterm and term-born infants at term-equivalent age (Fig. 3).

Among all networks, the default mode network (DMN) demonstrated the most pronounced developmental trajectory, exhibiting the largest difference between preterm and term-born infants (Fig. 3) and the steepest slope in maturation rate (Fig. 4). This rapid growth was also reflected in topographical changes, particularly in the frontal and temporal clusters, in both cross-sectional samples (Fig. 1) and a smaller longitudinal sample (Fig. 2). Although previous studies have shown postnatal development of the DMN (Doria et al., 2010; Sylvester et al., 2022), our findings reveal a more complete interconnection between discrete DMN clusters than reported in earlier studies (Eyre et al., 2021; Myers et al., 2024; Smyser et al., 2010).

The absence of a significant difference in slopes between the term and preterm maturational trajectories (Fig. 4) suggests that differences in functional development may already be established shortly after birth and are not substantially altered by the extra-uterine environment. However, further studies of fetal functional connectivity are needed to construct longitudinal curves for term-born infants over the same time span, enabling more direct comparisons between cohorts.

### Implications for early biomarkers

Precision functional mapping techniques have traditionally relied on large volumes of individual fMRI data, often collected over several hours (Gordon et al., 2017). Here, we present an alternative approach for estimating individual-level RSN maps using shorter fMRI scans, making it feasible to track individual neurodevelopment during the first weeks of life. When combined with other biomarkers collected later on, this approach could support the early identification of atypical developmental trajectories and inform personalized interventions.

Building on this, our subject-level analyses in the preterm cohort revealed both ongoing maturation and a significant reduction in functional connectivity strength across RSNs. These findings, in combination with recent studies linking structural MRI abnormalities (Pagnozzi et al., 2025; Trimarco et al., 2024) and diminished functional connectivity (Cyr et al., 2025) to motor outcomes in preterm infants, underscore the potential of individualized RSN maps as valuable tools for assessing early brain development and the potential impact of prematurity and other early-life exposures.

Given its rapid postnatal maturation and the pronounced differences observed between term and preterm infants, the DMN is a strong candidate biomarker for tracking early brain development. The fact that individual differences in the DMN are greater than in primary sensory networks, likely reflecting its later developmental trajectory, may also indicate greater inter-subject variability (as seen in the interquartile range, Fig. 3). This makes it especially promising for studying behavioral outcomes in early life, particularly as differences in DMN connectivity have already been linked to neurodevelopmental disorders like autism spectrum disorder (ASD) (Haghighat et al., 2022).

### Methodological considerations

While the longitudinal sample in this study only included 15 infants, the data provide valuable insights into individual developmental trajectories during this critical period of early brain maturation. Future work with larger longitudinal cohorts could better capture inter-individual variability and help refine our current estimates of developmental trajectories. Similarly, although a direct comparison between the longitudinal trajectories of preterm and term-born infants was not feasible within the current dataset, such an analysis would complement the cross-sectional findings (Fig. 4) and offer a more comprehensive picture of developmental differences between cohorts.

The current sample was also not ideally suited to examine the effects of gestational age as a continuous variable. In particular, the birth ages of subjects with term-equivalent scans (Fig. 3) were not evenly distributed, limiting our ability to assess the dose-dependent effects of prematurity. This remains an important direction for future research, as prior studies have shown significant differences in brain structure, functional connectivity, and long-term outcomes between infants born very preterm and those born moderate- or late-preterm (Dubois et al., 2021; Lemola et al., 2017; Padilla et al., 2015; Romeo et al., 2020).

Another potential source of bias in our study relates to head motion. A recent study (Ryan et al., 2023) has found a small but significant association between gestational age and sleep patterns in preterm infants. Furthermore, growing evidence suggests that head motion during fMRI is not merely a nuisance variable, but is linked to underlying neurophysiological states such as sleep and arousal (Blumberg et al., 2020; Denisova, 2019; Mitra et al., 2017). As a result, conventional motion correction techniques such as frame scrubbing may inadvertently bias the BOLD time series toward specific physiological states, and in this context, toward particular gestational ages. To reduce this risk, we opted to retain a contiguous block of data with the lowest motion, rather than relying on traditional scrubbing approaches.

## Conclusions

In this study, we applied a precision functional mapping approach to characterize individual resting-state network (RSN) patterns in preterm and term-born infants. This individualized method enabled detailed tracking of RSN development using short fMRI scans, supporting its potential for early biomarker identification. We found decreased functional connectivity in preterm infants, particularly within higher-order networks such as the default mode network (DMN), which also demonstrated the most rapid postnatal maturation. Although overall maturation rates were similar across groups, persistent differences in connectivity suggest that RSN disruptions related to prematurity may emerge shortly after birth. These findings highlight the value of subject-level mapping for detecting early developmental alterations and informing future longitudinal studies.

